# Effects of SGLT2 Ablation or Inhibition on Corticosterone Secretion in High-Fat-Fed Mice: Exploring a Nexus with Cytokine Levels

**DOI:** 10.1101/2024.04.18.590099

**Authors:** Niki F. Brisnovali, Isabelle Franco, Amira Abdelgawwad, Hio Lam Phoebe Tsou, Thong Huy Cao, Antonio Riva, Guy A. Rutter, Elina Akalestou

## Abstract

Despite recent therapeutic advances, achieving optimal glycaemic control remains a challenge in managing Type 2 Diabetes (T2D). Sodium-glucose co-transporter type 2 (SGLT2) inhibitors have emerged as effective treatments by promoting urinary glucose excretion. However, the full scope of their mechanisms extends beyond glycaemic control. At present, their immunometabolic effects remain elusive. To investigate the effects of SGLT2 inhibition or deletion, we compared the metabolic and immune phenotype between high fat diet-fed control, chronically dapagliflozin-treated mice and total-body SGLT2/*Slc5a2* knockout mice. SGLT2 null mice exhibited superior glucose tolerance and insulin sensitivity compared to control or dapagliflozin-treated mice, independent of glycosuria and body weight. Moreover, SGLT2 null mice demonstrated physiological regulation of corticosterone secretion, with lowered morning levels compared to control mice. Systemic cytokine profiling also unveiled significant alterations in inflammatory mediators, particularly interleukin 6 (IL-6). Furthermore, unbiased proteomic analysis demonstrated downregulation of acute-phase proteins and upregulation of glutathione-related proteins, suggesting a role in the modulation of antioxidant responses. Conversely, IL-6 increased SGLT2 expression in kidney HK2 cells suggesting a role for cytokines in the effects of hyperglycemia. Collectively, our study elucidates a potential interplay between SGLT2 activity, immune modulation, and metabolic homeostasis.

**Graphical Abstract:** 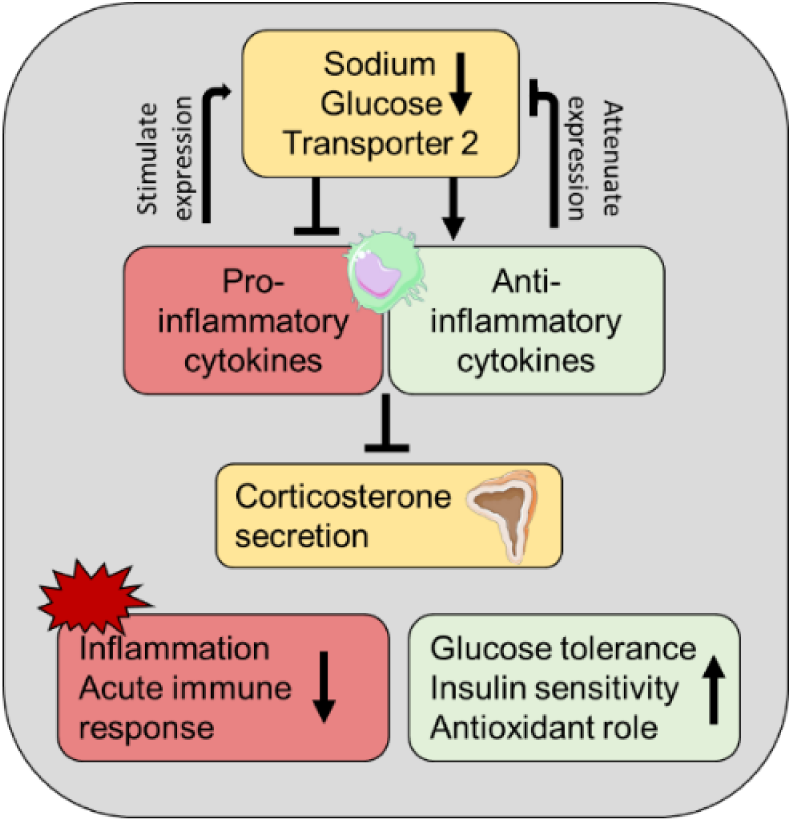

**Article Highlights:**

- The role of Sodium-glucose co-transporter type 2 (SGLT2) in immunity regulation remains elusive, despite extensive research in SGLT2 inhibitors.
- We sought to discern the effects of SGLT2 inhibition or deletion on metabolic and immune profiles in high-fat-fed mice, focussing on corticosterone regulation and cytokine alterations.
- SGLT2 null mice exhibit enhanced insulin sensitivity, alongside physiologically regulated corticosterone levels and significant alterations in inflammatory cytokines, and we identified changes in protein expression suggestive of antioxidant modulation.
- Our findings emphasize the interplay between immune responses and metabolic regulation mediated by SGLT2 activity.

---

The number of people living with diabetes in the UK reached a staggering 5 million in 2023, 90% of whom were affected by Type 2 Diabetes (T2D) (1). The National Diabetes Audit (NDA) 2021-2022 quarterly report for England revealed that glucose control deteriorated in patients with T2D, with only 35% of patients meeting the treatment targets for haemoglobin A1c (HbA1c) (2). Worldwide, the number of cases is predicted to reach 600 million by 2045 (3).

Despite significant pharmacological advances in recent years (4), there is still an unmet need to develop a more effective approach to tackling the pathophysiology of this disease.

Inhibition of the sodium-glucose co-transporter type 2 (SGLT2) has emerged as an effective means of improving glycaemic control in patients with T2D (5). SGLT2 is primarily expressed in the kidney cortex and inhibitors, known as gliflozins, block SGLT2 activity in the luminal membrane of the renal early proximal tubule to reduce glucose reabsorption by 50–60% (5). By lowering the renal glucose threshold to promote glucose excretion in the urine, glycaemia is lowered in an insulin-independent way (5).

SGLT2 inhibitors have been proven to lower HbA1c levels, increase high-density lipoprotein cholesterol and improve blood pressure, cardiovascular health and kidney disease (5). Despite their beneficial effects, their exact mechanisms of action are not fully understood, and may extend beyond renal glucose reabsorption. In particular, there is growing evidence for a broader, pleiotropic effect of SGLT2 inhibitors on immune regulation (6, 7), including inhibition of T-cell activation and proliferation (8) as well as modulation of macrophage M2 polarization (9). These findings hint at an intriguing immunometabolic role for SGLT2.

We have recently explored the role of SGLT2 (encoded by *Slc5a2*) in the metabolic responses to high fat diet and metabolic surgery, deploying knockout mice and pharmacological inhibitors (10). The aim of the present study was to evaluate the role of the immune system in the observed effects of SGLT2 ablation or inhibition.

## Research Design and Methods

### Animals

All animal procedures were approved by the British Home Office under the UK Animal (Scientific Procedures) Act 1986 with approval from the local ethical committee (Animal Welfare and Ethics Review Board, AWERB), at the Central Biological Services (CBS) unit at the Hammersmith Campus of Imperial College London. Adult male C57BL/6J mice (Envigo, Huntingdon U.K.) were maintained under controlled temperature (21-23°C), and light (12:12 hr light-dark schedule, lights on at 0700).

Mice carrying null *Slc5a2* alleles (*SGLT2*^−/−^ or “SGLT2 null^“^) lacking exon 1 were generated by CRISPR/Cas9-mediated recombination, as previously described (10). *11β-HSD1^−/−^*mice, also lacking exon one, were generated similarly (11) on a C57BL/6J background. In both cases, heterozygous animals were inter-crossed to generate wild-type, heterozygous and homozygous littermates.

Male mice were initially used as a continuation of our previous study (10) to allow direct comparison of results. We also conducted a pilot study using male and female SGLT2 null mice and found no differences in cytokine regulation measurements between the two groups. From the age of 8 weeks C57BL/6J control and SGLT2 null mice were placed on a 58 kcal% Fat and Sucrose diet (D12331, Research Diet, New Brunswick, NJ) to induce obesity and diabetes. Group 1 (n=5) received 10mg/kg dapagliflozin dissolved in methylcellulose daily for 14 days, through oral gavage. Biopsies (kidneys, adipose tissue, plasma) were harvested from all mice in the fed state, and were snap frozen in −80°C.

### Glucose tolerance tests

Mice were fasted for 8hrs and given free access to water. Blood was sampled in EDTA coated tubes from the tail vein at 0, 5, 15, 30, 60 and 90 min after intraperitoneal glucose administration (1g/kg). For the measurement of Glucagon-like Peptide 1, glucose (3 g/kg body weight) was administered via oral gavage following overnight (total 16 h) fasting and aprotinin was added in the collection tubes. Blood glucose was measured with an automatic glucometer (Accuchek; Roche, Burgess Hill, UK).

### Insulin tolerance tests

Mice were fasted for 8 h (08:00) and given free access to water. At 15:00, human insulin (Actrapid, Novo Nordisk) (1.5 U/kg body weight) was administered via intraperitoneal injection. Blood was sampled as described in Glucose Tolerance tests.

### Urine glucose and creatinine measurement

Urine was collected from mice during glucose tolerance tests by placing them on a cage without bedding. Glucose was measured by Glucose Assay (abcam, USA) and creatinine was measured by Creatinine Assay (Crystal Chem, USA). Urinary glucose-to-creatinine ratio (UGCR) was calculated as (mg/mg) = [urine glucose (mg/dl)/ urine creatinine (mg/dl).

### Cell culture and treatments

HK2 cells were obtained from ATCC (Manassas, VA) and were cultured with Keratinocyte Serum Free Medium (K-SFM) supplemented with 0.05mg/ml BPE and 5ng/ml EGF (Invitrogen), without added glucose. Cells were seeded at a density of × 10^5^ per well and treated for 72 hrs with glucose (30mM), IL-6 (1nM), leptin (5nM), FGF21 (10nM), IL-1β (1nM) or TGF1β (400pM) (Abcam, USA). A total of 4–10 different replicates were used for each condition.

NCI-H295R cells were obtained from ATCC (Manassas, VA) and cultured in DMEM:F12 medium, supplemented with 0.00625mg/ml insulin, 0.00625mg/ml transferrin, 6.25ng/ml selenium, 1.25mg/ml bovine serum albumin, 0.00535mg/ml linoleic acid and 2.5% Nu-Serum (Corning, USA). Cells were seeded at a density of × 10^5^ per well and treated for 24 hrs with 100ng/ml IL-6 (1nM), leptin (5nM), FGF21 (10nM), IL-1β (1nM), TGF1β (400pM) or IL-10 (1nM) (Abcam, USA). A total of 4–10 different replicates were used for each condition.

### RNA isolation, cDNA synthesis and quantitative polymerase chain reaction (qPCR)

RNA was extracted from cell and tissue samples by resuspending them in TRIzol reagent (Ambion, Life Technologies, Cat #349907) and chloroform (Sigma-Aldrich, Cat# SHBL1580). Total RNA isolation was conducted using the TRIzol™ Plus RNA Purification Kit (Applied Biosystems, Thermofisher) according to the manufacturer’s protocol. The High-Capacity cDNA Reverse Transcription kit (Applied Biosystems, Thermofisher) was used for cDNA synthesis. Master mix for qPCR was made using SYBR green (Applied Biosystems, ThermoFisher), RNase Samples were analysed using 7500 Fast Real-Time PCR system (Applied Biosystems, Serial #275010654). All data were normalised against *β-actin* as the housekeeping gene and all ΔCt values were calculated using the average from dual replicates. The analytical method was using 2^-ΔΔCt^. The primers used are shown in Table 1.

**Table 1:**
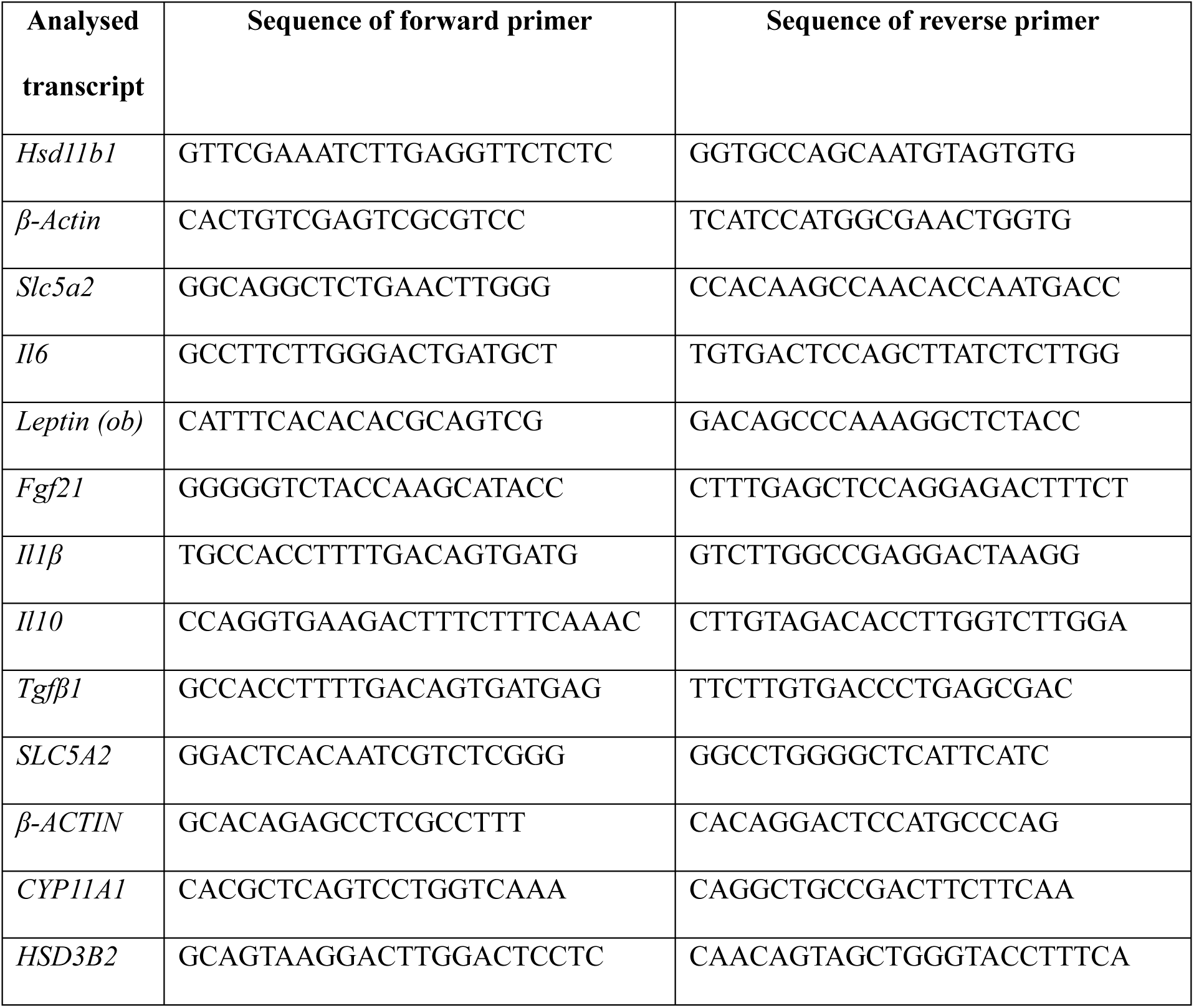
Primers used for the quantitative detection of transcripts normalized to β-actin.

*Enzyme-linked immunosorbent assays:* Blood plasma protein concentrations of corticosterone (Crystal Chem), GLP-1 (Crystal Chem) and TGFβ1 (Proteintech) were measured using their respective assay kits and processed according to manufacturer’s protocol.

### Multiplex cytokine quantification

Blood was collected at the end of treatment using a cardiac puncture. The samples were centrifuged at 10,000g for 8 minutes at 4°C to isolate the plasma. The blood plasma was stored at −20°C. Blood plasma cytokine concentrations were measured using a Luminex Mouse Discovery Assay (8-Plex) (LXSAMSM-07) according to manufacturer’s instructions. Standard curves were interpolated by 5-parameter logistic curves. All quantified values between sensitivity limits and lowest standard points were included in the analysis. All values below quantification limits were also included in the analysis as sensitivity or ‘half-minimum’ values.

### Proteomics analysis

Proteomics analysis was conducted by Seer Inc. (California, USA) through Liquid-Chromatography-Mass Spectrometry (LC-MS).

### Sample preparation for direct digest protocol

For direct digestion of neat plasma samples, proteins were denatured, reduced, alkylated and subjected to proteolytic digestion (Trypsin and Lys-C) for 3 hours at 37°C. Peptides were purified by solid phase extraction and yields were determined (Applied Biosystems, ThermoFisher).

### Sample preparation with the Proteograph™ XT Assay Protocol

Samples were processed with the Proteograph XT Assay (12). In brief, plasma proteins were quantitatively captured in nanoparticle (NP) associated protein coronas. Proteins were subsequently denatured, reduced, alkylated, and subjected to proteolytic digestion (Trypsin and LysC). Peptides were purified and yields were determined. Peptides were dried down overnight with a vacuum concentrator and reconstituted with a reconstitution buffer to a concentration 50ng/µL.

### Data-Independent Acquisition LC-MS/MS

For Data-Independent Acquisition (DIA), 8µl of reconstituted peptide mixture from each NP preparation was analyzed resulting in a constant 400ng mass MS injection between NP A and NP B samples. Each sample was analyzed with a Vanquish NEO nanoLC system coupled with a Orbitrap TM Astral TM (Applied Biosystems, ThermoFisher) mass spectrometer using a trap-and-elute configuration. First, the peptides were loaded onto an AcclaimTM PepMapTM 100 C18 (0.3mm ID x 5mm) trap column and then separated on a 50cm µPACTM analytical column (PharmaFluidics) at a flow rate of 1µl/min using a gradient of 5–25% solvent B (0.1% FA, 100% ACN) mixed into solvent A (0.1% FA, 100% water) over 22 min, resulting in a 30min total run time. The mass spectrometer was operated in DIA mode with MS1 scanning and MS2 precursor isolation windows between 380-980 m/z. MS1 scans were performed in the Orbitrap detector at 240,000 R every 0.6 seconds with a 5ms ion injection time or 500% AGC (500,000 ion) target. Two-hundred fixed window MS2 DIA scans were collected at the Astral detector per cycle with 3Th precursor isolation windows, 25% normalized collision energy, and 5 ms ion injection times with a 500% (50,000 ion) active gain control maximum. MS2 scans were collected from 150-2000m/z.

### Statistical analysis

Principal component analysis (PCA) was performed to determine if there was a good separation between three mouse groups (HFD vs. dapagliflozin treatment vs. SGLT2 null) using SIMCA version 14 (MSK Umetrics). The fold-change and adjusted p values in comparisons between three mouse groups were calculated using Limma R package with adjustment for Benjamini-Hochberg false discovery rate (FDR) of 0.05 to control multiple testing errors and then volcano plots were generated. Data were analyzed using GraphPad Prism 10 software. Comparisons between two groups were carried out using Mann-Whitney test. Group comparisons were analysed using a two-way ANOVA, given lack of missing measurements, with Geisser-Greenhouse correction, and Sidak’s multiple comparison test. The model included main effects and interaction, to be able to evaluate group differences at individual timepoints. All statistical tests were 2-tailed, and significance was set at alpha = 0.05. Errors signify ± Median with interquartile range.

### Data and resource availability

All data generated during this study are included in the published article (and its online supplementary files).

## Results

### SGLT2 null mice display improved glucose tolerance compared to dapagliflozin-treated mice independent of glycosuria levels

Whole body SGLT2 knockout mice and wild type (WT) littermate controls were exposed to 12 weeks of High Fat Diet (HFD). Subsequently, WT mice were treated with either dapagliflozin (10mg/kg) or control vehicle (methyl cellulose) via oral gavage daily for 10 days. In order to confirm deletion, levels of SGLT2/*Slc5a2* were measured in kidney cortex biopsies in all mice (Fig. 1a). Of note, *Slc5a2* tended to be elevated in dapagliflozin-treated compared to WT mice. Glucose to creatinine measurement ratio in mice urine following a 3g/kg glucose load showed no significant difference between the two intervention groups (Fig. 1b).

**Figure 1:**
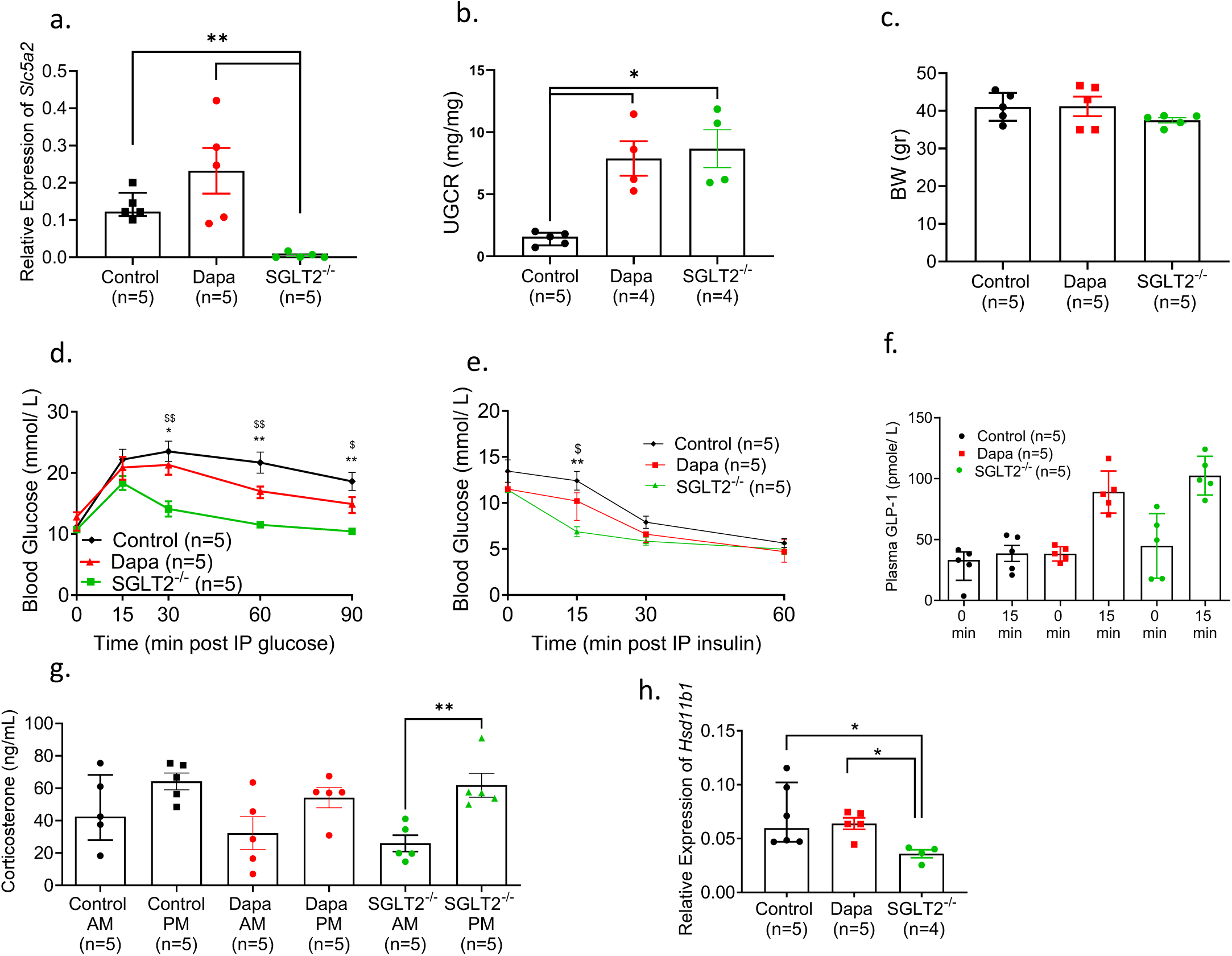
(a) Gene expression of SGLT2/*Slc5a2* relative to *β-actin* (*Actb*) in kidney of Wild Type High Fat Diet (HFD) fed mice following 4wk methyl cellulose or dapagliflozin oral gavage treatment (10mg/kg) and HFD SGLT2 null obese mice. (b) Urinary Glucose to Creatinine Ratio post IPGTT (1g/kg, 8hr fasting). (c) Body weight. (d) IPGTT (1g/kg, 8hr fasting) $ WT vs. SGLT2−/−,* WT+Dapa vs. SGLT2−/−, (e) ITT (1.5IU/kg, 6 hr fasting), $ WT vs. SGLT2−/−,* WT+Dapa vs. SGLT2−/− (f) Glucagon-like peptide 1 (GLP-1) levels following oral GTT (3g/kg, 16hr fasting). (g) Plasma corticosterone concentration in the morning [0800-0900] (AM) and evening [1700-1800] (PM). (h) Gene expression of 11β Hydroxysteroid dehydrogenase 1 (11β-HSD1)/ *Hsd11b1*, relative to *β-actin* in the subcutaneous adipose tissue *p<0.05, **p<0.001

No significant weight loss was observed in either group (Fig. 1c). SGLT2 null mice tended to have a lower body weight (37.5 ± 1.6g) than the dapagliflozin- and vehicle-treated mice (Av. 41.2 ± 5.8g and 41.0 ± 3.8g, p=0.10 and 0.93 respectively). During intraperitoneal glucose tolerance tests (IPGTT, 1g/kg) SGLT2 null mice displayed significantly improved glucose clearance (blood glucose total AUC 1175 ± 57.4mmol/ L 0-90min) when compared to control WT (blood glucose total AUC 1876 ± 121.5 mmol/ L 0-90min) and dapagliflozin-treated (blood glucose total AUC 1589 ± 52.1 mmol/ L 0-90min) WT mice (Fig. 1d). SGLT2 null mice also showed improved insulin tolerance (Fig. 1e).

To investigate whether these differences may reflect higher levels of the incretin glucagon-like peptide 1 (GLP-1), plasma levels of GLP-1 were measured after 16h of fasting and following oral gavage (3g/kg glucose). Compared to untreated animals, glucose-induced GLP-1 increases tended to be restored in SGLT2 null (p=0.06, 0 vs. 15min, n=5) and dapagliflozin-treated WT (p=0.06, n=5, 0 vs. 15min) mice versus HFD control (Fig. 1f).

### Corticosterone secretion and Hsd11b1 expression in SGLT2 null mice is inhibited

To further explore the mechanisms behind the significantly improved glucose clearance and insulin tolerance in SGLT2 null mice, and more minor effects on these parameters in dapagliflozin-treated WT animals, we measured the levels of corticosterone in the morning (AM) and evening (PM) in plasma (Fig 1g). Obese WT mice demonstrated high levels of corticosterone in the morning followed by a marginal postmeridian increase, consistent with studies reporting increased levels of corticosterone (cortisol in humans) in obesity (13). In contrast, SGLT2 null mice demonstrated a more physiological secretion pattern, reflecting tendences towards both lowered AM and increased PM corticosterone levels (Fig. 1g). Correspondingly, expression of the corticosterone-activating enzyme 11β-Hydroxysteroid dehydrogenase type 1 (11βHSD1/*Hsd11b1)* in subcutaneous adipose tissue (Fig. 1h), was significantly reduced in SGLT2 null mice (p=0.015), but unaffected in dapagliflozin treated animals.

### SGLT2/Slc5a2 expression and glycosuria are unaffected in 11βHSD1 null mice

To explore the possibility of a bidirectional relationship between SGLT2 and 11βHSD1 expression, we used total body 11βHSD1/*Hsd11b1* knockout mice (11) and assessed SGLT2*/Slc5a2* expression and function after 12 weeks on HFD. No significant differences in weight were observed between the two groups (Fig. 2a). No difference in SGLT2*/Slc5a2* expression was observed in the kidney cortex (Fig. 2b) or the glucose to creatinine ratio (Fig 2c).

**Figure 2:**
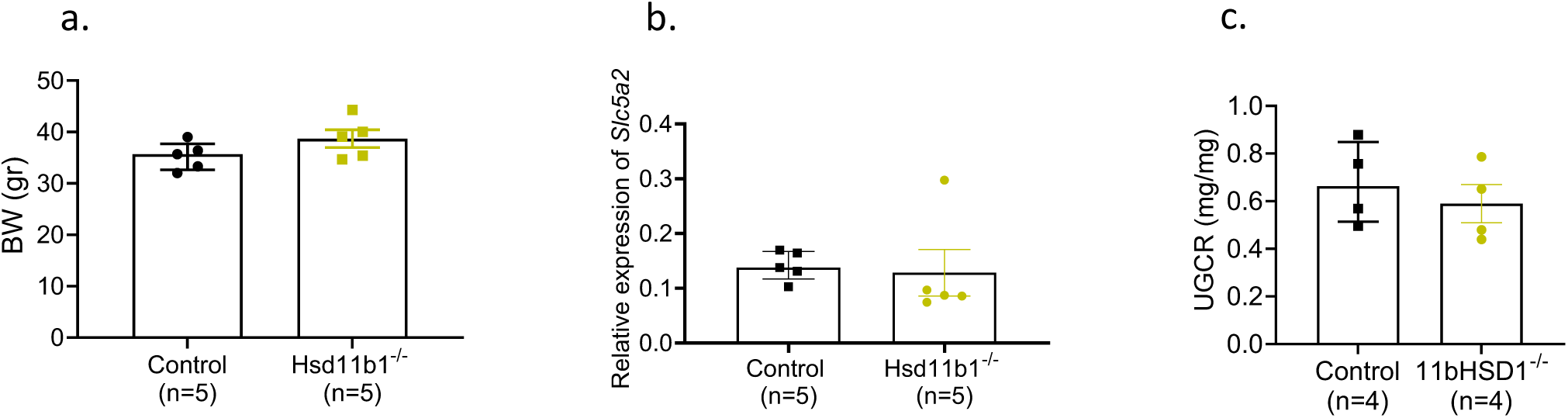
(a) Body weight of Wild Type High Fat Diet (HFD) induced obese WT and 11β-HSD1)/ *Hsd11b1* null obese mice. (b) Gene expression of SGLT2/*Slc5a2* relative to *β-actin* in kidney. (c) Urinary Glucose to Creatinine Ratio post IPGTT (3g/kg)

### SGLT2 null mice demonstrate altered circulating cytokine levels in plasma and tissue

Following our finding that corticosterone secretion is normalised in SGLT2 null mice, we sought next to investigate whether the effects of SGLT2 deletion or inhibition on glucose homeostasis and corticosterone levels may involve altered levels of cytokines. The pro-inflammatory cytokine IL-6, clearly detectable in WT HFD mice, was significantly lowered (p=0.079) in SGLT2 null animals (Fig. 3a). In contrast, levels of the anti-inflammatory cytokines IL-10 and FGF21 were significantly increased compared to WT mice (p=0.01 and p=0.02 respectively) (Fig. 3d, e). Of note, IFNg, TNFa and IL-1b were always below detection range. To verify these findings, we measured the gene expression of the most prominent cytokines in subcutaneous adipose tissue biopsies of the same SGLT2 null and WT mice. Both *Il6* and *leptin* (*ob*) genes were downregulated in SGLT2 null mice, while *Il10* and *Tgfb1* were upregulated (Fig. 4) versus WT mice.

**Figure 3:**
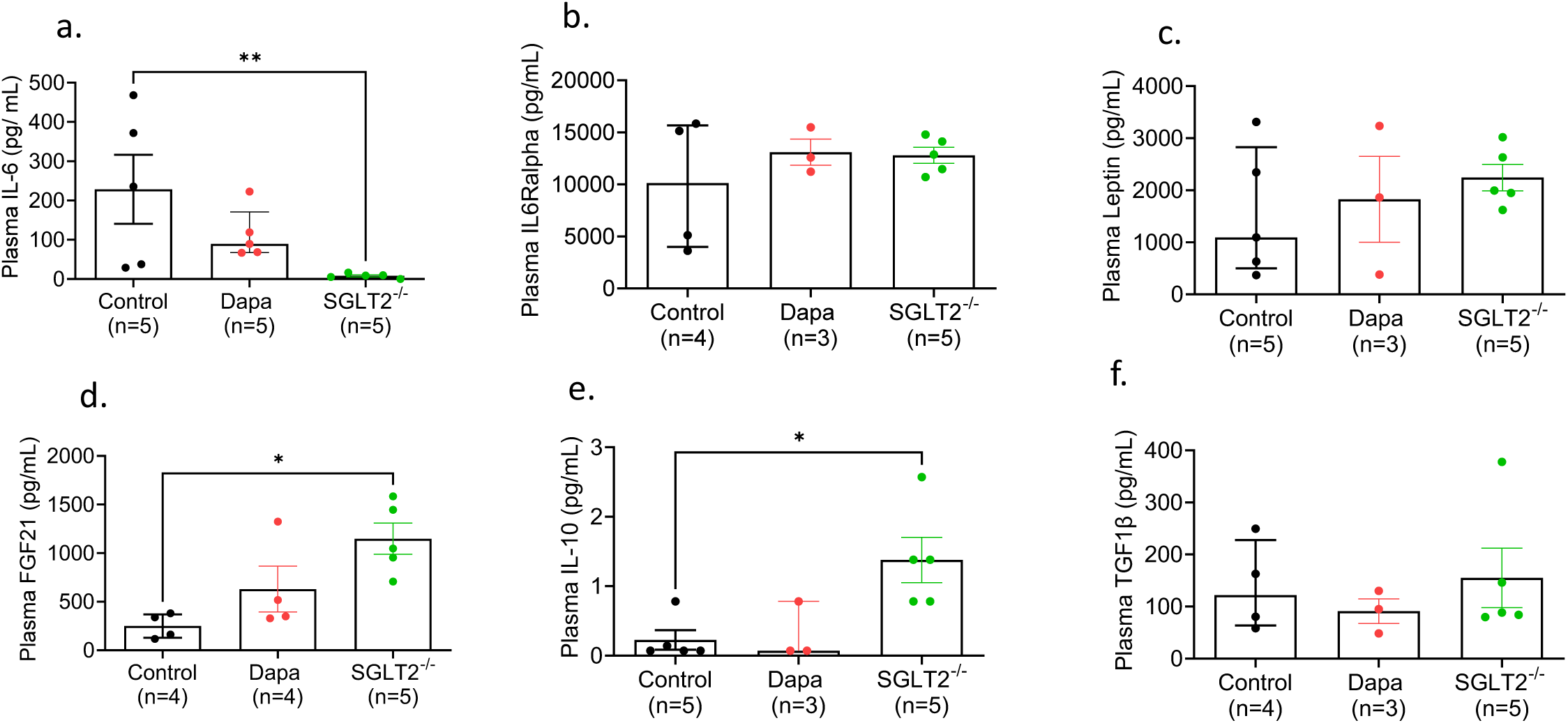
(a) Plasma concentration of IL-6. (b) Plasma concentration of IL-6 Receptor alpha (c) Plasma concentration of leptin. (d) Plasma concentration of FGF21 (e) Plasma concentration of IL-10. (f) Plasma concentration of TGFβ in Wild Type High Fat Diet (HFD) induced obese WT following 4wk methyl cellulose or Dapagliflozin treatment (10mg/kg) and SGLT2 null obese mice. *p<0.05, **p<0.001

**Figure 4:**
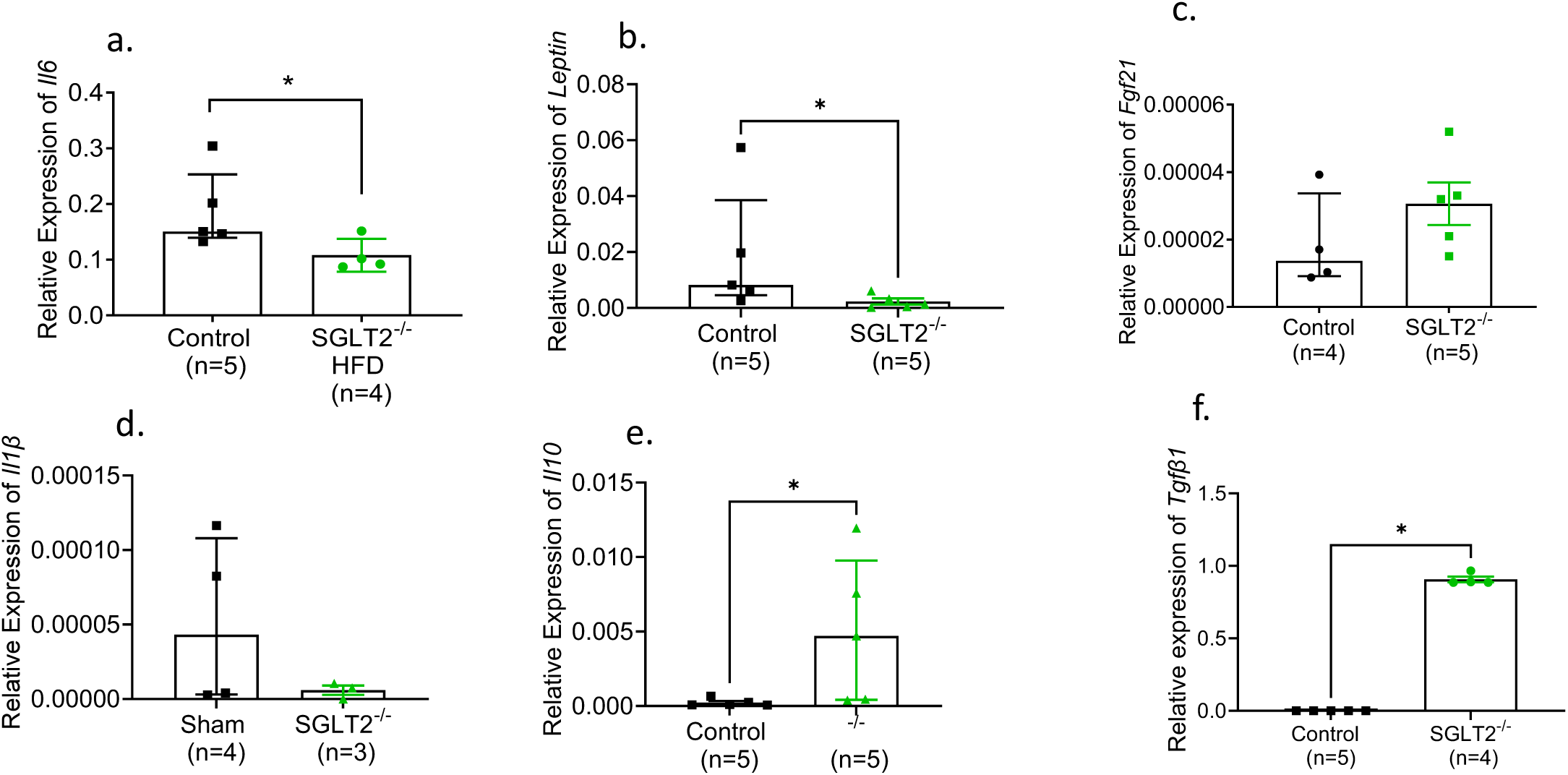
(a) Gene expression of *Il6* (b) *Leptin* (c) *Fgf21* (d) *Il1β* (e) *Il10* (f) *Tgfβ1* relative to *β-actin* in the subcutaneous adipose tissue of Wild Type High Fat Diet (HFD) induced obese WT and SGLT2 null mice *p<0.05

### Circulating levels of proteins involved in inflammation were downregulated in both dapagliflozin-treated and SGLT2 null mice

Plasma from the three mouse groups was further analysed using mass spectrometry. Principal component analysis showed clear difference between the groups (Fig. 5a). We quantified 5660 plasma proteins, 127 of which were downregulated and 252 were upregulated in dapagliflozin-treated mice, when compared to WT control (Fig. 5b). In the SGLT2 null mice, 57 proteins were downregulated and 233 were upregulated, when compared to WT control (Fig. 5c). Of these proteins, 18 were down-regulated and 92 were upregulated in both dapagliflozin-treated and SGLT2 null mice. We narrowed the targets by focusing on the proteins with the highest fold-change (>1.5-fold change) and lowest p-value threshold (p<0.05). Functional enrichment analysis was then performed using Database for Annotation, Visualization and Integrated Discovery (DAVID) (14, 15) (https://david.ncifcrf.gov/). Protein interaction networks were drawn using the Search Tool for the Retrieval of Interacting Genes/Proteins (STRING) database (16) (https://string-db.org/). Both dapagliflozin-treated and SGLT2 null mice demonstrated significant downregulation of 6 proteins involved in acute liver response to possible immune challenge and immune system processes: Fibrinogen-Like Protein 1, Amyloid P Component, Haptoglobin, Serum Amyloid A1, Serum Amyloid A2, Serum Amyloid A3 (Fig. 5d). Additionally, 4 proteins also involved in vessel morphogenesis were downregulated in dapagliflozin-treated mice: Thrombospondin 1, Fms-related tyrosine kinase 1, Hepatocyte growth factor, Angiopoietin 1 (Fig. 5d).

**Figure 5:**
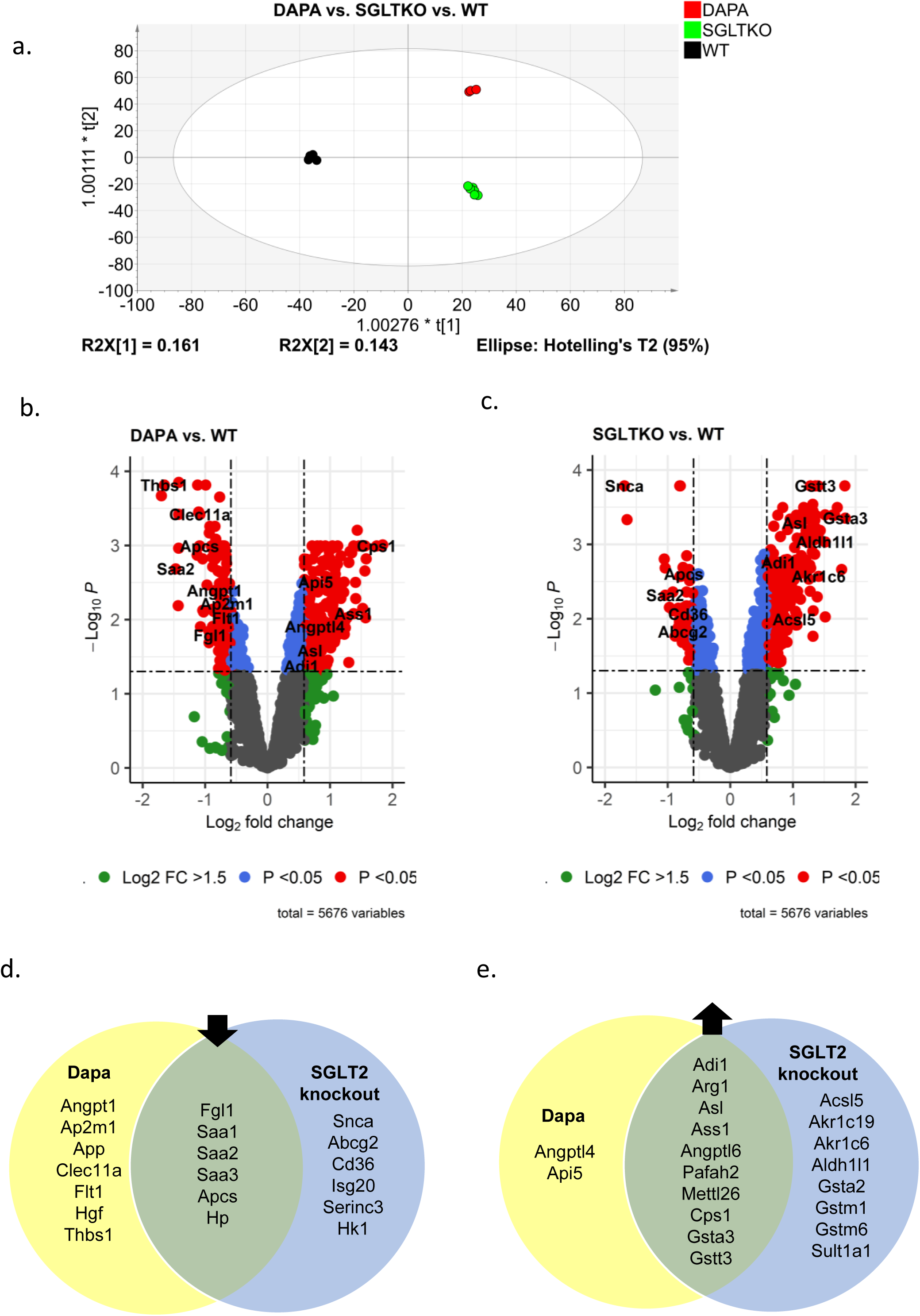
(a) Principal component analysis (PCA) plot of three groups: (HFD) induced obese WT following 4wk methyl cellulose (black, n=6), Dapagliflozin treatment (10mg/kg) (red, n=3), and SGLT2 null obese mice (green, n=6) (b) Volcano plot showing differentially abundant plasma proteins when comparing Dapa-treated mice vs. WT. (c) Volcano plot showing differentially abundant plasma proteins when comparing SGLT2 null mice vs. WT. Genes associated with differentially abundant proteins have been annotated. (d) Venn diagram displaying immune response-related proteins downregulated in Dapa-treated (yellow) or SGLT2 null (blue) mice, or both (green). (e) Immune response-related proteins upregulated in Dapa-treated (yellow) or SGLT2 null (blue) mice, or both (green)

### SGLT2 null mice exhibit upregulated expression of proteins involved in glutathione production

Of the 92 proteins found to be upregulated in both dapagliflozin-treated and SGLT2 null mice, four (Argininosuccinate synthase 1, Arginase 1, Argininosuccinate lyase, Carbamoyl-phosphate synthase 1) are enzymes involved in the urea cycle (Fig. 5e), as well as cellular responses to glucagon. Importantly, three proteins involved in “glutathione metabolic process” were upregulated in both groups, compared to WT (Glutathione S-transferase alpha 3, Glutathione S-transferase theta 3, Glutathione S-transferase Mu 3) (Fig. 5e), while SGLT2 null mice displayed significantly upregulated expression of 11 additional glutathione-regulating proteins.

### Cytokines directly alter SGLT2/SLC5A2 expression in human kidney cells

To determine whether increased SGLT2/*SLC5A2* expression may involve inflammatory mediators, we treated human kidney cells (HK2) with a selection of the cytokines flagged during the *in vivo* experiments above (Figs 3, 4). The expression in HK2 cells of the cognate receptor for each cytokine was confirmed by RT-qPCR. The cell line was first treated with high glucose (30mM) and revealed a dose-responsive expression of *SLC5A2* (Fig. 6a). The pro-inflammatory cytokines IL-6 and leptin, corresponding to those with reduced gene expression (and/or plasma levels) in SGLT2 null mice (Fig 4a, b), also strongly induced *SLC5A2* (Fig 6b, c), compared to untreated controls. Conversely, cells treated with the anti-inflammatory cytokines FGF-21, IL-10 and TGFβ, corresponding to those with increased gene expression (and/or plasma levels) in SGLT2 null mice (Fig 4c, e, f), displayed significantly lower *SLC5A2* expression (Fig 6d, e, f),

**Figure 6:**
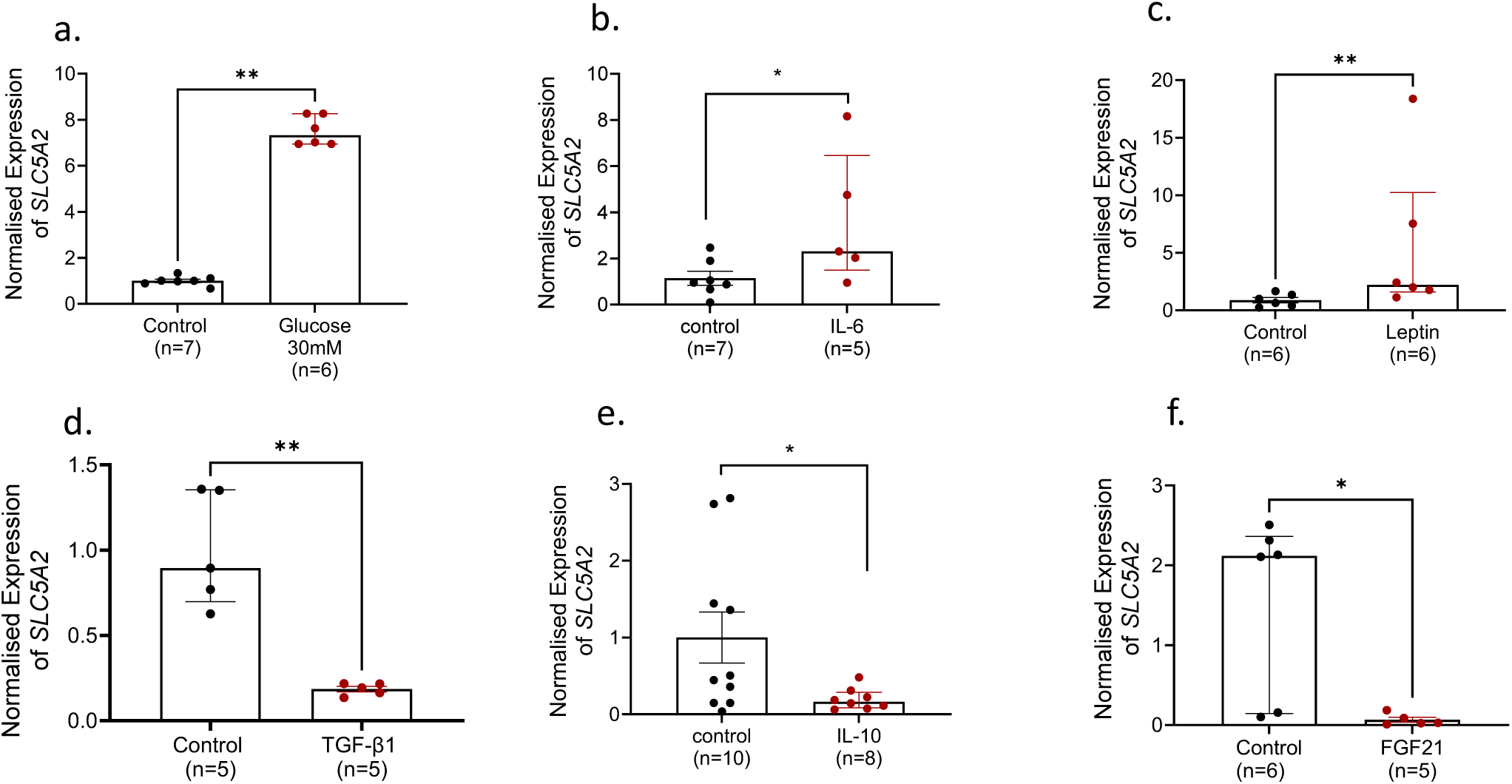
(a) Normalized gene expression of SGLT2/*SLC5A2* relative to β-actin in the human kidney HK2 cell lysates following 96hr treatment with (a) glucose (30mM) (b) IL-6 (1nM) (c) Leptin (5nM) (d) TGF1β (400pM) (e) IL-1β (1nM) (f) FGF21 (10nM) *p<0.05, **p<0.001

### Pro-inflammatory cytokines increase the expression of key cortisol-regulating enzymes

To explore the possibility that altered cytokine levels (Figs 3, 4) may contribute to the apparent normalisation of corticosterone concentration after SGLT2 deletion (Fig. 1g, h) we treated the human adrenal cortex cell line H295R with the previously-identified cytokines (Fig 3, 4, 5). After 24 h treatment, the pro-inflammatory cytokine IL-6 (and, to a lesser extent leptin) increased the expression of the cortisol synthesis enzymes encoded by *CYP11A1* and *HSD3B2.* The anti-inflammatory cytokines IL-1β1, TGFβ1, IL-10 and FGF21 had no effect (Fig. 7a, b).

**Figure 7:**
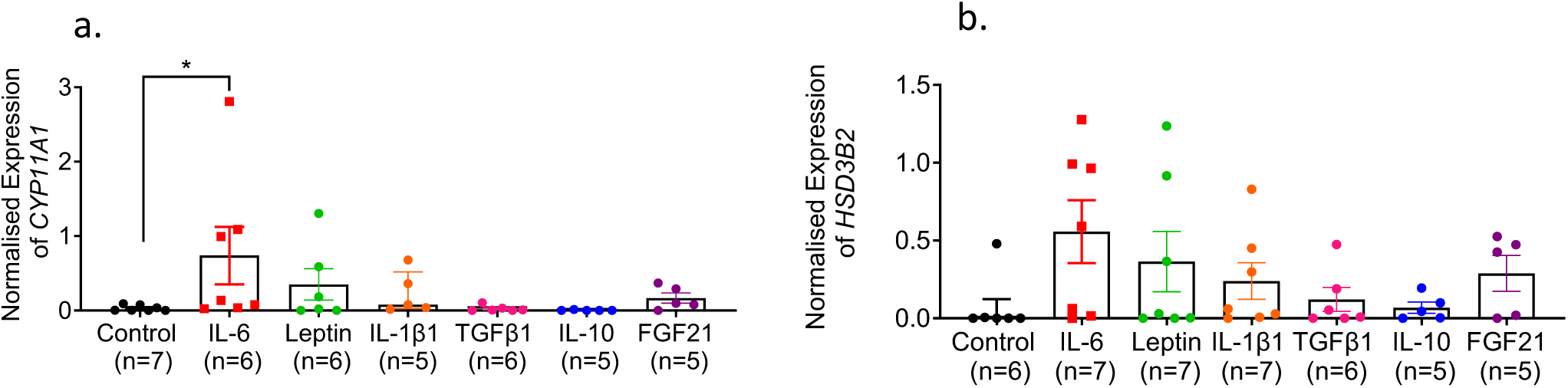
Normalized gene expression of (a) *CYP11A1* and (b) *HSD3B2* relative to *β-actin* in the human adrenal gland H295R cells following 24hr treatment with (i) untreated medium (ii) IL-6 (1nM) (iii) Leptin (5nM) (iv) IL-1β (1nM) (v) TGF1β (400pM) (vi) IL-10 (1nM) (vii) FGF21 (10nM) *p<0.05

## Discussion

We show here that SGLT2 null mice on a high fat diet have improved glucose tolerance and insulin sensitivity, compared to dapagliflozin-treated mice. This effect appears to be independent of glycosuria levels and may be due to a reduction in cytokine and consequent corticosterone secretion.

Corticosterone (cortisol in humans) is a steroid hormone secreted primarily by the adrenal glands. Its main functions include increasing blood glucose through gluconeogenesis and suppression of the immune response. However, excess cortisol and high levels of its activating enzyme 11βHSD1 are linked to profound metabolic disturbances resulting in insulin resistance and dyslipidaemia (13), as well as increased production of inflammatory cytokines (17). Obesity has often been associated with cortisol dysregulation and excess secretion, further contributing to the state of low-grade inflammation (13). We show that complete ablation of SGLT2 restores a normal “physiological” fluctuation (11) in corticosterone in HFD mice, an effect which also tended to be observed, albeit to a lesser extent, after dapagliflozin-treatment. Consistent with these findings, the SGLT2 inhibitor empagliflozin has been reported to improve chronic hypercortisolism-induced abnormal myocardial structure and cardiac function in mice (18). Similarly, tofogliflozin is reported to reduce cortisol levels in patients with T2D (19). The observation that 11βHSD1 null mice have normal levels of SGLT2 gene expression and glycosuria (Fig. 2), indicates that the reduced corticosterone in SGLT2 null mice is likely an outcome, rather than a part in a feedback loop (Fig. 1g, h).

We also observed significantly lower levels of in the pro-inflammatory cytokine IL-6 in SGLT2 null mice, compared to the two WT groups. Although dapagliflozin-treated mice did not achieve significantly lower IL-6 concentrations, as previously described (20, 21), they did display lower levels compared to vehicle-treated control mice. However, plasma levels of soluble IL-6 receptor alpha (sIL-6Ra) remained unchanged. Soluble IL-6 receptor alpha is commonly found in the peripheral circulation and can capture free IL-6, expanding the range of cells capable to mount pro-inflammatory responses to IL-6 (22). Of note, leptin levels remained comparable to control mice in both dapagliflozin-treated and SGLT2 null mice. Previous studies have shown that pharmacological inhibition of SGLT2 reduces leptin levels (23). However, this effect was linked to SGLT2-inhibitor therapy-induced weight loss, an effect that is largely absent in our model and may explain the overall lack of change in cytokine circulation in dapagliflozin-treated mice. We note that it has been previously shown that HFD SGLT2 null mice did not display significant weight loss, compared to WT controls (24). The anti-inflammatory cytokines FGF-21 and IL-10, known for their role in improving insulin sensitivity (25) and inhibiting inflammation (26), were present at higher circulating levels in SGLT2 null vs control WT mice, while TGFβ1 concentration was unchanged.

We further evaluated these findings in SGLT2 null mice, by measuring the gene expression of cytokines in subcutaneous adipose tissue. Our plasma results of IL-6, FGF21 and IL-10 were replicated in adipose tissue gene expression. Additionally, *leptin* gene expression was significantly reduced in SGLT2 null mice, while *TGFβ1* was increased, even though their levels were unchanged when measured in plasma. This could indicate a more tissular role, or it may be related to the lack of association between gene expression and plasma cytokine levels. Leptin is primarily produced in adipose tissue and high levels are tightly associated with insulin resistance and inflammation (27). The role of TGFβ1 is less clear. It can be produced, secreted and stored in the extracellular matrix, inactive and without functional consequences until tissue remodelling processes are induced (28). In the adipose tissue, TGFβ1 has been shown to be upregulated in obese mice (29). However, its role in the regulation of inflammatory responses depends on the differentiation stage of adipocytes (30, 31), while TGFβ receptor 1 has been shown to regulate progenitors that promote browning of white fat (32). As SGLT2 null mice did not demonstrate significantly different body, compared to WT controls, it is likely that the increase in TGFβ1 is linked to the latter effect.

Upon analysis of plasma proteomes in the three mice groups, we found that both dapagliflozin-treated and SGLT2 null mice had reduced levels of acute phase proteins, which are produced by the liver in response to pro-inflammatory stimuli, such as IL-6 or other pro-inflammatory cytokines. Fibrinogen-Like Protein 1 (FGL1) has a well-known tumorigenic role but is also considered an immune checkpoint due to its role in modulating T-cell function, when bound with the immune inhibitory receptor Lymphocyte-activation gene 3 (LAG3) (33). Haptoglobin is an acute-phase protein which is found elevated in several inflammatory diseases (34). Similarly, Serum Amyloid A proteins have a critical role in control and possibly propagation of an initial acute phase response (35). As expected, both interventions resulted in the upregulation of enzymes involved in the urea cycle, linked to the acute contraction in plasma volume caused by SGLT2-inhibition (36). Of note, glutathione-regulating proteins were also found to be upregulated in both groups, compared to control, especially in SGLT2 null mice. This indirect antioxidant role has recently been reported in SGLT2 inhibitors and it is hypothesised to be due to their ability to reduce high glucose-induced oxidative stress (37, 38).

Providing evidence of a role for altered cytokine levels in the observed normalisation of corticosterone fluctuations in SGLT2 null mice, treatment of adrenal gland cells with IL-6 upregulated *CYP11A1* and *HSD3B2*, in line with previous findings (39). An upward trend was also observed following treatment with leptin (40) and IL-1β (41), while IL-10 and TGFβ1 had no effect, again mirroring earlier results (42, 43). Interestingly, FGF21 appeared to cause a slight increase in the expression of the two enzymes, an effect previously described in rodents as a part of a glucocorticoid to FGF21 feed-forward loop (44). Nevertheless, a direct demonstration of a role for inflammatory cytokines in mediating the effects of SGLT2 deletion (or inhibition) in the living animal will require inactivation of the corresponding cytokine or receptor genes, or immunostasis, which lie beyond the scope of the present study.

We also demonstrate that pro-inflammatory cytokines IL-6 and leptin caused an increase in SGLT2 gene expression, while anti-inflammatory cytokines FGF21, IL-10 and TGFβ1 caused a decrease, compared to untreated control. This raises the possibility of a bidirectional relationship between SGLT2 and cytokine regulation.

This study has limitations, importantly the low sample sizes in the dapagliflozin-treated mouse group in proteomic analysis. This is a result of high volumes of plasma required for cytokine measurement and mass spectrophotometry. Moreover, reduced function of SGLT2, as a result of altered gene expression in HK2 cells, was not confirmed. Future studies should include further interrogation of SGLT2 expression and functional regulation through glucose uptake detection *in vitro*, following treatment with different agents. Additionally, proteomics analysis of multiple key immunometabolic tissues and cell populations, such as liver, spleen and macrophages should be performed.

In summary, we found that genetic ablation of SGLT2 exerted more profound effects than pharmacological inhibition. Our data overall confirm previous studies reporting an effect of SGLT2 inhibition on cytokine secretion (45–47), which may be attributed to a number of mechanisms such as macrophage polarisation (9), T-cell suppression (8) or resolution of low-grade inflammation through glycosuria (23). However, SGLT2 null mice demonstrated a significantly improved metabolic phenotype, compared to SGLT2 inhibitor-treated mice. This effect was likely driven by a significant reduction in pro-inflammatory cytokines, as well as an increase in anti-inflammatory cytokines, which subsequently led to physiological levels of corticosterone secretion.

As glycosuria, GLP-1 and body weight levels were similar between the two interventions, this observation opens questions on the physiological role of SGLT2 in immune response, the modulatory role of hyperglycaemia, and the temporal relationship between its metabolic effects and the anti-inflammatory effects reported in this study. Answering these questions could unravel a new mechanism of action for SGLT2 inhibitors, and potentially a new avenue for the development of for immunometabolic treatments.

## Author contributions

**Niki F. Brisnovali, Isabelle Franco, Amira Abdelgawwad:** Methodology, Investigation, Formal analysis, Visualization. **Hio Lam Phoebe Tsou:** Investigation, Formal analysis. **Thong Huy Cao:** Software, Formal analysis, Writing - Review & Editing, Visualization. **Antonio Riva:** Conceptualization, Formal analysis, Writing - Review & Editing, Visualization. **Guy Rutter:** Conceptualization, Resources, Writing - Review & Editing **Elina Akalestou:** Conceptualization, Validation, Investigation, Formal analysis, Funding acquisition, Writing - Original Draft, Visualization, Supervision.

## Guarantor Statement

Elina Akalestou is the guarantor of this work and, as such, had full access to all the data in the study and takes responsibility for the integrity of the data and the accuracy of the data analysis.

## Conflict of interest

G.A.R. has received grant funding from Sun Pharmaceuticals Inc. for whom he is a consultant. No other potential conflicts of interest relevant to this article were reported.

## Funding

N.F., I.F., A.A. and E.A. were supported by a British Heart Foundation Accelerator Grant (AA/18/3/34220) and a Project Grant from the Rosetrees Trust (M825). G.A.R. was supported by a Wellcome Trust Investigator Award (212625/Z/18/Z) UKRI-Medical Research Council (MRC) Programme grant (MR/R022259/1), an NIH-NIDDK project grant (R01DK135268) a CIHR-JDRF Team grant (CIHR-IRSC TDP-186358 and JDRF 4-SRA-2023-1182-S-N), CRCHUM start-up funds, and an Innovation Canada John R. Evans Leader Award (CFI 42649). A.R. and H.L.P.T. are supported by the Foundation for Liver Research (London, UK). T.H.C. was funded by the National Institute for Health and Care Research Leicester Biomedical Research Centre.

